# Genotyping of *Leptospira* spp. in wild rats leads to first time detection of *L. kirshneri* serovar Mozdok in Serbia

**DOI:** 10.1101/2024.01.11.575145

**Authors:** Vladimir Gajdov, Goran Jokic, Sara Savic, Marina Zekic, Tanja Blazic, Milica Rajkovic, Tamas Petrovic

## Abstract

This study aimed to investigate the prevalence and molecular characterization of *Leptospira* species in Belgrade, Serbia, an area where this disease is underexplored. Specifically, the study sought to employ molecular and multilocus sequence typing analyses to fill the gap in understanding the diversity and distribution of *Leptospira* species within the region. A comprehensive molecular analysis was conducted on kidney samples obtained from Norway rats (*Rattus novegicus*) in urban environments. The study utilized molecular diagnostic techniques including real-time PCR targeting the *lipL32* gene and performing sequence-based typing schemes utilizing *adk, icdA, lipL32, lipL41, rrs2* and *secY* genes. These methodologies were applied to ascertain the presence and characterize different *Leptospira* species and serotypes, respectively. The findings revealed the presence of two *Leptospira* species and three separate serotypes in the Belgrade area. Moreover, this study identified the presence of *L. kirschneri* serovar Mozdok in Serbia for the first time, a significant discovery previously undocumented in the region. This pioneering investigation sheds light on the molecular diversity and prevalence of *Leptospira* species in Serbia. The study underscores the importance of employing molecular typing methods to gain insights into the epidemiology and characterization of *Leptospira* species. These findings significantly contribute to both local and global perspectives on leptospirosis epidemiology, providing vital insights for the development of effective control strategies and interventions.

**Author summary:** In our recent study, we explored the presence and performed molecular typing of the *Leptospira* species, the bacteria responsible for leptospirosis, in wild rats in Serbia. This was the first time such a study was conducted in the region. Leptospirosis is a serious disease that affects both animals and humans, often transmitted through contact with water contaminated by infected animals. Our focus was on understanding which types of *Leptospira* were present in these animals. Excitingly, we discovered a particular strain of *Leptospira*, known as *L. kirshneri* serovar Mozdok, for the first time in Serbia. This finding is significant because it sheds light on the presence and spread of different *Leptospira* serovars in Serbia. It also raises awareness about the potential health risks associated with this serovar, which was previously unknown in the area. Our work fits into a broader context of disease surveillance and public health. By identifying the types of Leptospira present in a specific region, we can better understand the risks to public health and take steps to prevent and control the spread of leptospirosis. This discovery is not just important for scientists studying infectious diseases; it has real implications for public health officials, veterinarians, and anyone concerned with preventing and treating leptospirosis. Our findings highlight the need for ongoing monitoring of *Leptospira* in wildlife, to protect both animal and human health.

## 1. Introduction

Leptospirosis, a zoonotic disease caused by pathogenic spirochaetes of the genus *Leptospira*, is constantly present in some parts of the world and holds significant relevance in both veterinary and public health contexts due to its ability to cross over between humans, domestic animals, wildlife and even environment (water). Reported cases of leptospirosis are global with over one million cases annualy, leading to approximately 60,000 fatalities [1]. To date, a minimum of 64 distinct *Leptospira* species have been validated worldwide using the average nucleotide identity (ANI) values of their genomes. While rats are traditionally known as the primary reservoirs for pathogenic *Leptospira* species, there have been numerous reports on various vertebrate and invertebrate hosts as excreting this pathogen through their urine. Wild and domestic mammals [2,3], livestock [4,5], amphibians [6], reptiles [7] and bats [8] also appear to play significant roles in the spread of *Leptospira* sp. Human infections typically result from exposure to soil or water contaminated with *Leptospira*, mostly from the urine of reservoir animals [9]. Detecting *Leptospira* through traditional growth on media can be problematic due to their slow growth, making it impractical for timely diagnoses. To address this, molecular diagnostic methods, such as the real-time PCR of the *lipL32* gene, have been developed [10, 11]. PCR-based amplification of *secY* and *ompL1* genes using species-specific primers and probes has been used to identify *Leptospira* species directly from clinical samples. These assays can identify common pathogenic *Leptospira* species when combined with a *lipL32* assay, including *L. borgpetersenii, L. interrogans, L. kirschneri*, and *Leptospira noguchii* [12]. Furthermore, sequence-based typing schemes utilizing gene targets like 16S rRNA *rrs2, secY*, and *lfb1*, or *adk, icdA, lipL32, lipL41, rrs2 and secY* have been developed for *Leptospira* [13,14]. For example, a ∼435-bp fragment of the *secY* gene shows good phylogenetic discrimination between pathogenic *Leptospira* species. Sequence-based methods can also be applied directly to clinical samples to determine the infecting species and genotype, as well as investigate links between human and animal *Leptospira* infection [15]. In Serbia, the presence of pathogenic *Leptospira* sp. has been documented in various animals including small wild mammals [16], however most of the studies in Serbia have been focused on seroprevalence and seroepidemiological detection of antibodies in samples from cats [17], dogs [18], cattle and sheep [19] and humans [20]. To the best of our knowledge this is the first study to perform molecular and multilocus sequence typing analysis of *Leptospira* species in Serbia. Moreover, this study revealed the presence of *Leptospira kirshneri* serovar Mozdok in Serbia for the first time.

## 2. Results

All 344 samples were analyzed for the presence of pathogenic *Leptospira* species. In kidney tissues, *Leptospira* spp. was detected in a total of 103 out of 344 individuals (29.94 %, 95% CI: 25.15-35.09) upon amplification by qPCR (Table 4). A total of 27 out of 103 positive samples (with Ct values between 20 and 28) were used in this study. Among all samples, the BLASTn analysis indicated that 26 sequences were affiliated with the *L. interrogans*, and 1 sequence exhibited the closest resemblance to the *L. kirschneri* (with 100% identity). The calculated sequence similarity of our samples with a cutoff value of 95% performed with Biopython was in concordance with the BLASTn results and for some of the samples it was possible to determine the serovar. For the final and definite characterization of our samples we determined the allele profile using the MLST scheme 3 from the PubMLST (https://pubmlst.org/Leptospira) database [24]. The MLST analysis yielded the following results: 11 of our samples belong to *L. interrogans* serovar Copenhageni, 12 to *L. interrogans* serovar Icterohaemorrhagiae and one to *L. kirschneri* serovar Mozdok. For the rest 3 of our samples, we were only able to determine the taxonomy to the level of species (*L. interrogans*) due to lower sequence quality.

## 3. Discussion

There is a growing interest in the surveillance of *Leptospira* spp. hosts, and investigations into the prevalence of this pathogen in wild mammals across Europe are on the rise and the significance of rodents as reservoirs for various *Leptospira* serovars has been extensively explored worldwide with various results. It is well-established that wild rats (*Rattus* spp.) are the principal sources of *Leptospira* infection, particularly in urban and peri-domestic environments [25]. The brown rat is known as the primary host of *L. interrogans* related to the serogroup Icterohaemorrhagiae, which is responsible for the most severe forms of the disease in humans [26]. This study aimed to examine the circulating *Leptospira* strains in wild rats, utilizing qPCR for initial detection of pathogenic *Leptospira* and MLST analysis for molecular characterization. Our findings confirm that wild rats harbor different serovars of pathogenic *Leptospira* spp. which pose threat to both animal and public health, highlighting the importance of continuous monitoring the presence and diversity of these bacteria in wild animals. The identification of *L. interrogans* serovar Icterohaemorrhagiae and *L. interrogans* serovar Copenhageni aligns with studies from all over Europe: in Sicily the bacteria has been detected in stray dogs and cats [27]; In Sardinia authors have reported pathogenic *Leptospira* in hedgehogs, mustelids and wild rodents [28]; In Germany, researchers in one study reported that 6% of the tested animals (various small mammals) exhibited positive results for *L. kirschneri* and *L. interrogans* [29], while *L. interrogans* serovar Icterohaemorrhagiae has been reported in wild rats all over the world [25] which is not surprising given that it represents the most common serovar in animals and humans. Additionally, this study relied on the utilization of the *adk, icdA, lipL32, lipL41, rrs2* and *secY* partial genes as a means for molecular typing and differentiating *Leptospira* serovars. The results obtained using these genes align with those obtained from other MLST analyses. Although Leptospirosis has been the subject of numerous studies across various geographical regions, this present investigation in Serbia marks a significant contribution to the field. Prior research in Serbia had mainly focused on seroprevalence and seroepidemiological studies [16-20]. However, our study distinguishes itself as the first in Serbia to employ molecular and multilocus sequence typing analysis for *Leptospira* species. This unique approach has yielded in discovering the presence of *Leptospira kirshneri* serovar Mozdok in Serbia. This marks the first documented occurrence of this serovar in the country. Similar reports have been documented in Croatia (30). The comprehensive and systematic testing conducted in our study, which included various *Leptospira* genes, facilitated the detailed characterization of positive samples. The sequencing and BLASTn analysis unveiled a predominance of *L. interrogans* in our samples, reinforcing its role as a common pathogenic *Leptospira* species. Further analysis, including the calculation of sequence similarity and allele profiling using the PubMLST database, refined our understanding of the *Leptospira* strains present. Notably, our findings unveiled specific serovars, such as *L. interrogans* serovar Copenhageni and *L. interrogans* serovar Icterohaemorrhagiae, underscoring the diversity of *Leptospira* strains within the Belgrade region. The significance of our discovery of *Leptospira kirshneri* serovar Mozdok in Serbia extends beyond the confines of our study. This novel serovar presence has far-reaching implications for vaccine strategies and epidemiological studies in both human and veterinary epidemiology. The discovery of *Leptospira kirshneri* serovar Mozdok in Serbia introduces a new dimension to vaccine development strategies. Serovars play a crucial role in vaccine formulation, as they determine the specific *Leptospira* strains that the vaccine should target. The presence of a novel serovar implies the need for the inclusion of this serovar in regional or local vaccine formulations. Failure to account for the presence of this serovar could compromise the effectiveness of vaccines in protecting both human and animal populations. Consequently, our findings serve as a critical foundation for the adaptation of vaccine strategies to the unique epidemiological landscape of Serbia. The present vaccine strategies in Serbia include preparations for different animals which contain *L. interrogans* serovar Icterohaemorrhagiae, Canicola, Copenhageni and Bratislava, *L. kirshneri* serovar Grippotyphosa. Regarding leptospiros epidemiology, the identification of *L. kirshneri* serovar Mozdok opens doors to a more comprehensive understanding of the disease’s distribution and dynamics in the region. The serovar’s presence highlights the complexity of *Leptospira* populations in Serbia and warrants further investigation into its reservoir hosts and transmission dynamics. Epidemiological studies must now consider the unique characteristics of this serovar, as it may exhibit distinct patterns of host adaptation and disease transmission. Understanding the prevalence and distribution of this serovar is crucial for developing effective control measures, both in terms of prevention and treatment. Moreover, the discovery emphasizes the importance of continued surveillance and monitoring of *Leptospira* diversity in the region, as new serovars may continue to emerge over time. In conclusion, our study has provided valuable insights into the presence and diversity of *Leptospira* species in Serbia. The discovery of *L. kirshneri* serovar Mozdok serves as a pivotal point for advancing vaccine strategies and epidemiological research in the region. By adapting our approaches to the unique characteristics of this novel serovar, we can better address the challenges of leptospirosis and work towards more effective prevention and control measures for both human and veterinary health. Furthermore, the presence of *Leptospira kirshneri* serovar Mozdok opens new avenues for epidemiological research in Serbia. This novel serovar’s presence highlights the complexity of *Leptospira* populations in the. Further research is essential to unveil the full implications of this discovery and to refine our understanding of the epidemiological landscape in Serbia.

## 4. Materials and methods

### 4.1 Animal Collection

The research was conducted in accordance with ethical principles and was approved by the Ministry of Agriculture, Forestry and Water Management (Republic of Serbia) - Veterinary Directorate (No. 323-07-04943/2020-05/2, 29.05.2020 and 323-07-04155/2023-05/2, 16.05.2023). During 2020, 2021 and 2022, a total of 344 (186 female and 158 male) carcasses of Norway rats (*Rattus norvegicus*) were collected in the broad environs of Belgrade City. Carcasses were collected predominantly in their urban and suburban habitats. The largest number of individuals was collected after the implementation of control measures or the implementation of monitoring measures. The collected carcasses were kept in a freezer at -20 ◦C for a short time, until further processing. During autopsy, the kidneys were separated for further analysis and the morphological data, body weight and sex of the animals were recorded.

### 4.2 DNA extraction, molecular detection, sequencing and MLST analysis

DNA was extracted from the kidney using the Quick-DNA MiniPrep kit (Zymo Research, Australia, Cat. no. D3024), according to manufacturers’ instructions. Due to validate the extraction processes and all downstream steps, nuclease-free water and DNA extracted from *Leptospira* positive samples were used as positive and negative controls, respectively. DNA extracted from each sample was stored at −20 °C until downstream use. To distinguish between pathogenic and non-pathogenic *Leptospira*, we performed qPCR targeting the *lipL32* partial target genes. Specifically, we used primers LipL32F (5’-GGA TCC GTG TAG AAA GAA TGT CGG-3’) and LipL32R (5’-GTC ACC ATC ATC ATC ATC GTC C-3’) to amplify a 101 bp fragment of the *lipL32* gene, which was detected by the probe LipL32P (6-carboxyfluorescein [FAM]-5’-ATG CCT GAC CAA ATC GCC AAA GCT GCG AAA-3’-Black Hole Quencher 1 [BHQ1]) [10]. An internal control, represented by exogenous DNA added before the extraction phase, representing simultaneously the extraction and PCR amplification control (qPCR Extraction Control RED, Meridian Bioscience, UK) was also included. The qPCR was carried out in a 12 μL reaction mixture containing 3 μL of *Leptospira* spp. genomic DNA, 0.5 μL (concentration of 20 pmol/μL) of forward and reverse primer and probe and 5 μL (concentration of 10pmol/μL) of FastGene 2x PROBE Universal (Nippon Genetics, Germany) and 2.5 μL of PCR water. All reactions were conducted in duplicates using a 7500 Fast Real-Time PCR System (Applied Biosystems, ThermoFisher, USA) with the following conditions: initial denaturation at 95°C for 2 min, followed by 45 cycles of denaturation at 95°C for 20 s, and annealing/elongation at 65°C for 50 s. Each PCR test included a negative control (DNA extracted from water) and a positive control (DNA extracted *Leptospira* spp. positive samples). Among the positive samples obtained through qPCR, only those with threshold cycle (Ct) values lower than or equal to 30 underwent further analysis. Specifically, 27 kidney samples and 27 *Leptospira* isolates were subjected to PCR using a set of primers amplifying *adk, icdA, lipL32, lipL41, rrs2* and *secY* partial genes (Table 1) [14]. PCR reagents and their volumes, as well as PCR cycling conditions are shown in Table 2 and Table 3, respectively. The PCR products were visualized by electrophoresis on a 1.5% agarose gel and examined under UV transillumination.

**Table 1:**
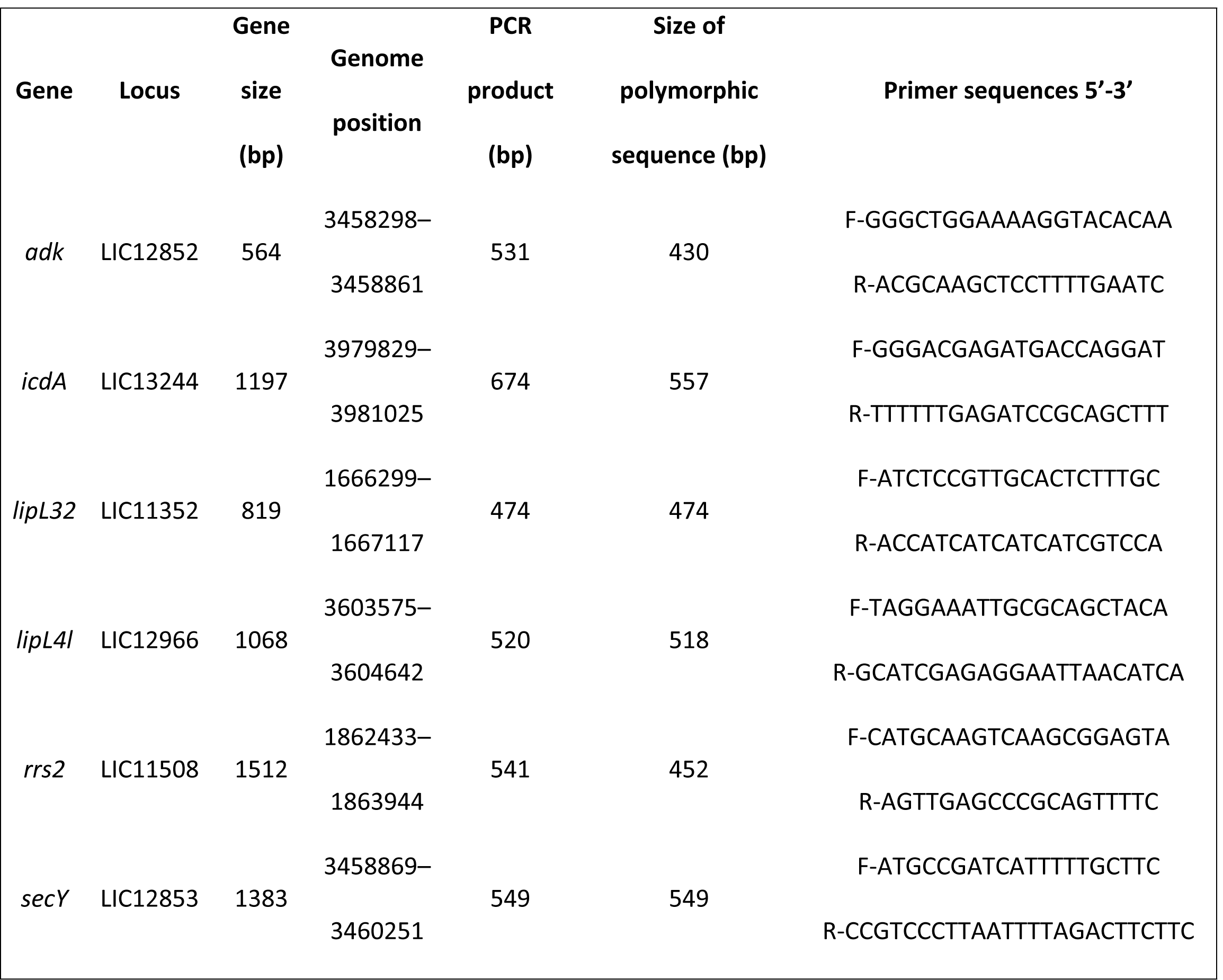
Details of gene loci and the corresponding primer sequences used for MLST Analysis.

**Table 2:** Reagents and Volumes.

| Reagent | Volume per<br>Reaction - $\mu\text{L}$ |
| --- | --- |
| DNA Template (*Leptospira*<br>DNA) | 3 |
| Forward Primer | 1 (concentration of<br>20 pmol/$\mu\text{L}$) |
| Reverse Primer | 1 (concentration of<br>20 pmol/$\mu\text{L}$) |
| HotStarTaq Master Mix | 12.5 |
| Sterile Water | 7.5 |
| Total Reaction Volume | 25 |

**Table 3:**
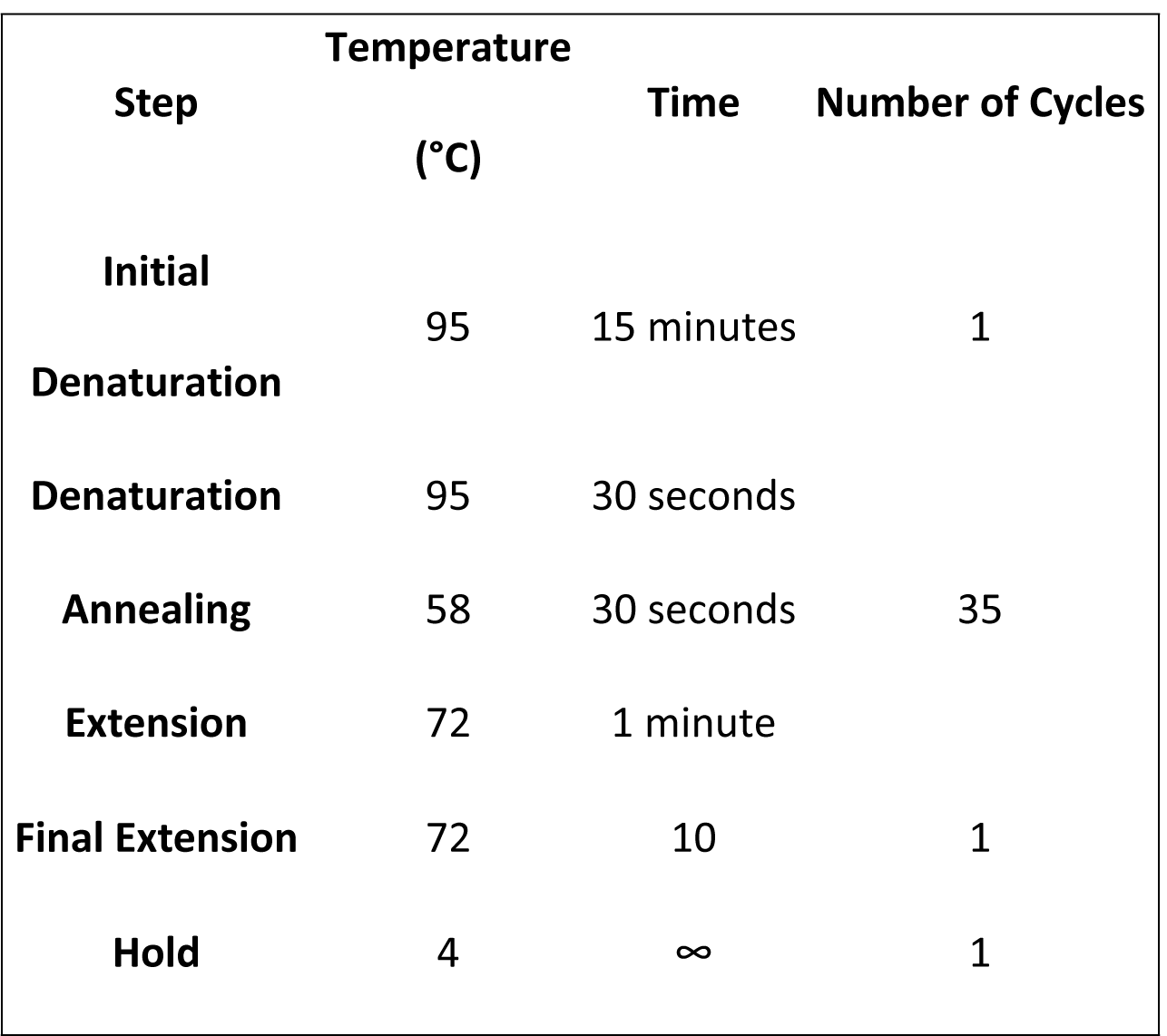
PCR Cycling Conditions.

**Table 4:**
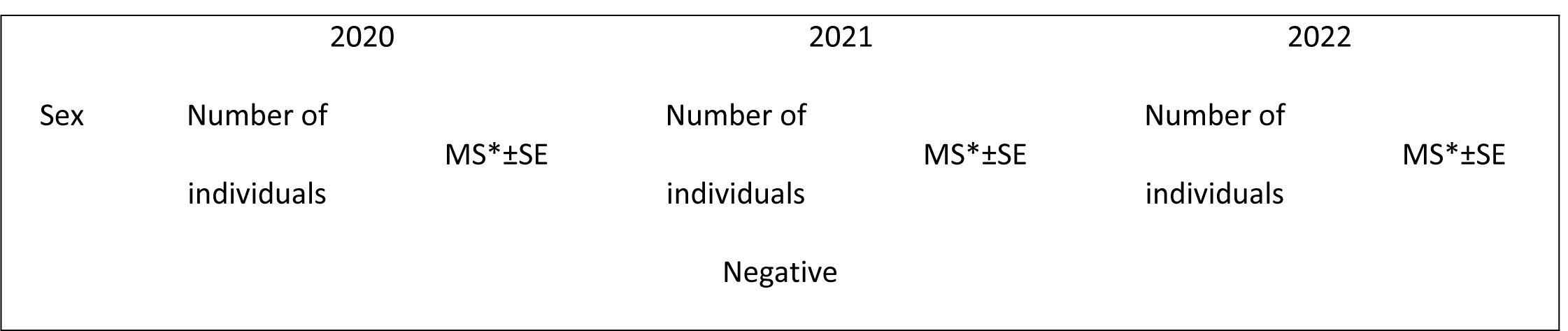
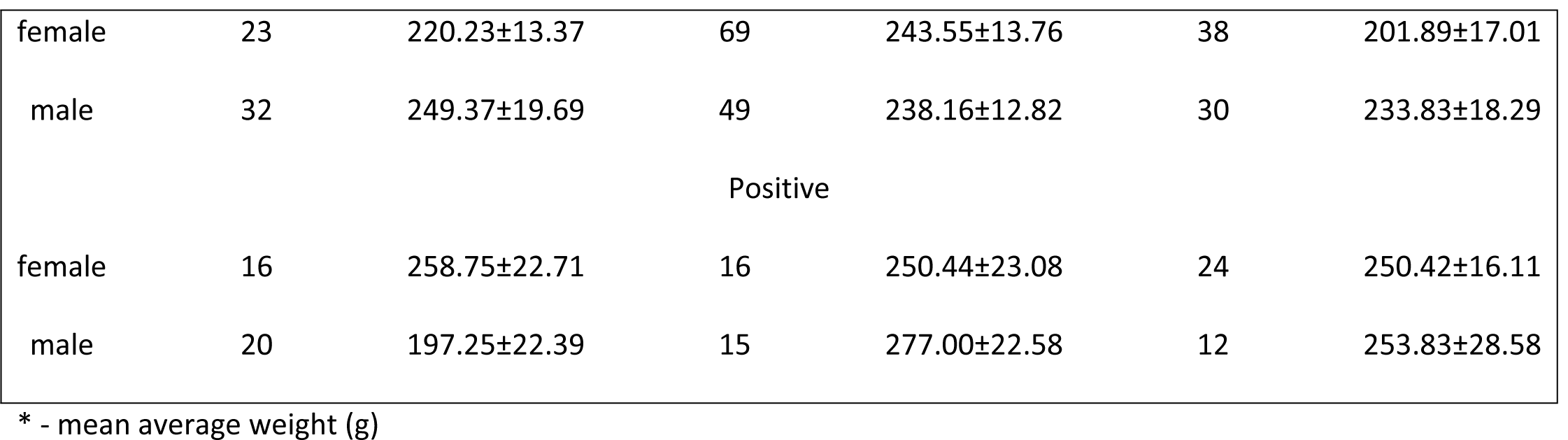
The presence of *Leptospira* spp. in Norway rat kidney tissues, collected in the period 2020.-2022. in Belgrade, Serbias.

We purified (GeneJET PCR Purification Kit, ThermoFisher Scientific, USA, cat. no. K0702) and sent all positive amplicons for genes listed in Table 2 to Macrogen Europe for Sanger sequencing. Sequences were analyzed and edited using the Staden package [21]. Consensus sequence validation was performed against a custom *Leptospira* database using nucleotide blast (BLASTn) [22] Each allele and the allelic profiles (adk-icdA-lipL32-lipL41-*rrs2*-secY) were submitted to the *Leptospira* database [23] (http://pubmlst.org/Leptospira, accessed in October 2023) for ST assignment. Sequence similarity of our samples was performed with a custom reference database using Biopython [24]. All sequences were submitted to NCBI’s GenBank under the following accession numbers: **<u>OR920389</u> - <u>OR920523</u>** for *adk, icdA, lipL32, LipL41* and *secY*, while for *rrs2* **<u>OR912477</u>-<u>OR912503</u>**.

### 4.3 Statistical analysis

Mean prevalence and confidence intervals (95% CI) for *Leptospira* spp. were determined using the Clopper and Pearson method. An alpha value <0.05 was considered the threshold for statistical significance.

## Ethical Approval statement

Ethics review and approval for this study were obtained from the Ministry of Agriculture, Forestry and Water Management (Republic of Serbia) - Veterinary Directorate (No. 323-07-04943/2020-05/2, 29.05.2020 and 323-07-04155/2023-05/2, 16.05.2023).

